# Inter-regional delays fluctuate in the human cerebral cortex

**DOI:** 10.1101/2022.06.01.494224

**Authors:** Joon-Young Moon, Kathrin Müsch, Charles E. Schroeder, Taufik A. Valiante, Christopher J. Honey

## Abstract

The flow of information between cortical regions depends on the excitability at each site, which is reflected in fluctuating field potentials. It remains uncertain how global changes in field potentials affect the latency and strength of cortico-cortical couplings. Therefore, we measured changes in oscillations and inter-regional couplings by recording intracranially from the human cerebral cortex. As participants listened to an auditory narrative, global increases in low-frequency (4-14 Hz) power were associated with stronger and more delayed inter-regional couplings. Conversely, increases in broadband high-frequency power were associated with weaker coupling and zero lag. In network oscillator models, these changes in cortico-cortical latency can be generated by varying the effective influence of inter-regional projections relative to intra-regional dynamics. Altogether, low-frequency oscillations appear to modulate information flow across the human cerebral cortex, as they covary with the timing of peak excitability between regions, and this process may be regulated by nonspecific ascending projections.

## Introduction

Although each patch of the cerebral cortex is anatomically connected to many others, the couplings between regions continually shift as we act, think and perceive. Electrophysiological studies have indicated that fluctuations in cortical excitability, as indexed by field potential oscillations, can regulate the flow of information along specific brain pathways ^1–3^ and can affect upcoming behavior ^4–7^. However, little is known about how the millisecond-scale couplings between brain regions are related to global oscillatory fluctuations in the human cerebral cortex. Therefore, we set out to measure changes in inter-regional coupling recorded in intracranial electrophysiological measurements from the human brain. We asked: how does the strength and latency of couplings between regions change with oscillatory power? We focused on the delays (latencies) between field potentials in different regions because if fluctuation in the electrical potential in one brain area leads that in another area, then this affects the likelihood that spikes in one area will elicit spikes in the other ^8–11^. More generally, patterns of zero-lag and nonzero-lag coupling are associated with distinct functional states ^12–17^.

Intracranial recordings from the human brain have revealed a large-scale pattern of latencies, in which parietal regions often lead temporal regions ^10,18,19^. Some of these latency patterns reflect the occurrence of 4-14 Hz waves traveling along the human cortical surface ^19^. Pioneering studies in the cerebral cortex of the cat indicated that the delays between brain regions became longer when the animals switched from a task-period to a rest period, and that this shift in delays was accompanied by an increase in alpha-band (10Hz) oscillations ^12^. Therefore, we hypothesized that, super-imposed on a default pattern of parietal-to-temporal flow, the inter-regional delays would continually fluctuate in the human brain, becoming longer when low-frequency oscillatory processes were stronger. We tested this hypothesis by measuring the inter-regional delays in intracranial recordings from human participants listening to minutes of natural narrative speech.

We found that increases in low-frequency power (4-14 Hz oscillations, LF power) were associated with longer conduction latencies between regions and stronger inter-regional correlations overall. Thus, increases in low-frequency power, both locally (at each recording electrode) and globally (averaged across the lateral cerebral cortex), were associated with a shift from zero-lag coupling to nonzero-lag (delayed) corticocortical coupling. In a small number of sites, the time-varying changes in inter-regional delays could be reliably elicited by presenting a time-varying auditory stimulus; however, the bulk of the latency fluctuations we observed did not show reliable locking to external drivers, and appeared to be endogenously controlled.

To gain insight into the neurophysiological processes that may underlie these changes in inter-regional delay, we modeled the dynamics as a system of phase-and-amplitude coupled oscillators. Our formal model accounted for the data in the following way: when the dynamical influence between regions is increased (e.g. via diffuse neuromodulation or thalamocortical signaling that changes the coupling gain) the amplitude of low-frequency oscillations increases, as does the magnitude and the latency of inter-regional correlations.

These data reveal an organization principle between the mesoscopic and macroscopic dynamics in the human brain: more desynchronized neuronal populations couple to other populations more weakly and with shorter delays, while more synchronized cortical populations couple to other regions more strongly and with longer delays. We consider this set of findings in terms of prior work proposing a role for ascending neuromodulatory projections which regulate arousal, as well as the balance between top-down and bottom-up information flow ^20–23^.

## Results

### Latencies fluctuate

Inter-channel latencies were identified using the cross-correlations of the raw voltage traces measured from subdural ECoG electrodes. We analyzed data from 10 participants as they listened to auditory narratives (2 presentations of a 7min 19s stimulus, Table S1 for participants’ information). For each channel pair, we computed the cross-correlation of the voltage signal in 2-second sliding windows (Fig. 1*A*). Within each time window, we defined the time lag, τ, as the time-delay associated with maximal inter-electrode correlation (Fig. 1*B*, see Methods). The cross-correlograms and τ values varied over time, as illustrated by an example pair in auditory pathway along the superior temporal gyrus (Fig. 1*C*, mean τ = 5.14 +/-6.30 ms).

**Fig. 1.**
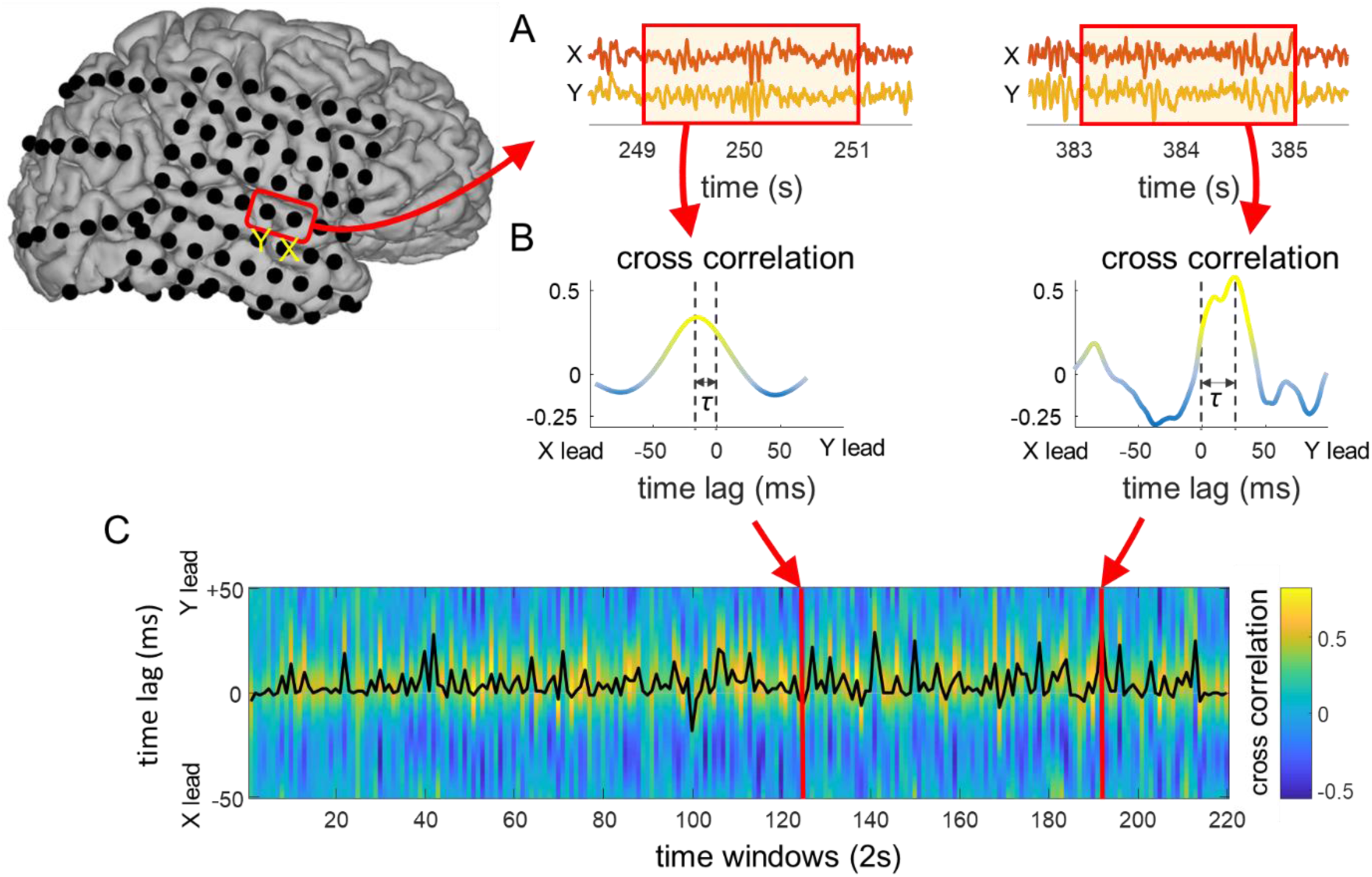
Schematic of latency analysis. We presented a 7-minute auditory narrative twice to each participant. **(*A*)** Moving time windows were applied to the pair of chosen channels. **(*B*)** Within each time window, we measured the cross correlation between the pair. The latency τ was defined as the time lag yielding the maximum cross-correlation between the channels. **(*C*)** Cross correlation (blue to yellow colors) and latency τ (black line) of this channel pair are illustrated for all time windows. The latency τ fluctuates throughout the time course.

### Inter-channel latencies increase with low-frequency oscillatory amplitude

Our initial investigation of electrodes in the auditory pathway identified many sites in which latencies were longer during increase of low-frequency (4-14 Hz) power and were shorter during increases of high-frequency broadband (65+ Hz) power (Fig. 2*B*). For example, for a neighboring electrode-pair in the superior temporal gyrus (STG, Fig. 2), alpha-band (10 Hz) power in those electrodes within a 2s time window was positively correlated with the inter-electrode latency for that window (Spearman ρ = 0.55, p<<0.01), while broadband power was negatively correlated with the latency (Spearman ρ = -0.36, p <<0.01). Therefore, we set out to characterize the relationship between low-frequency (LF) oscillations strength and inter-regional delays across the lateral cerebral cortex.

**Fig. 2.**
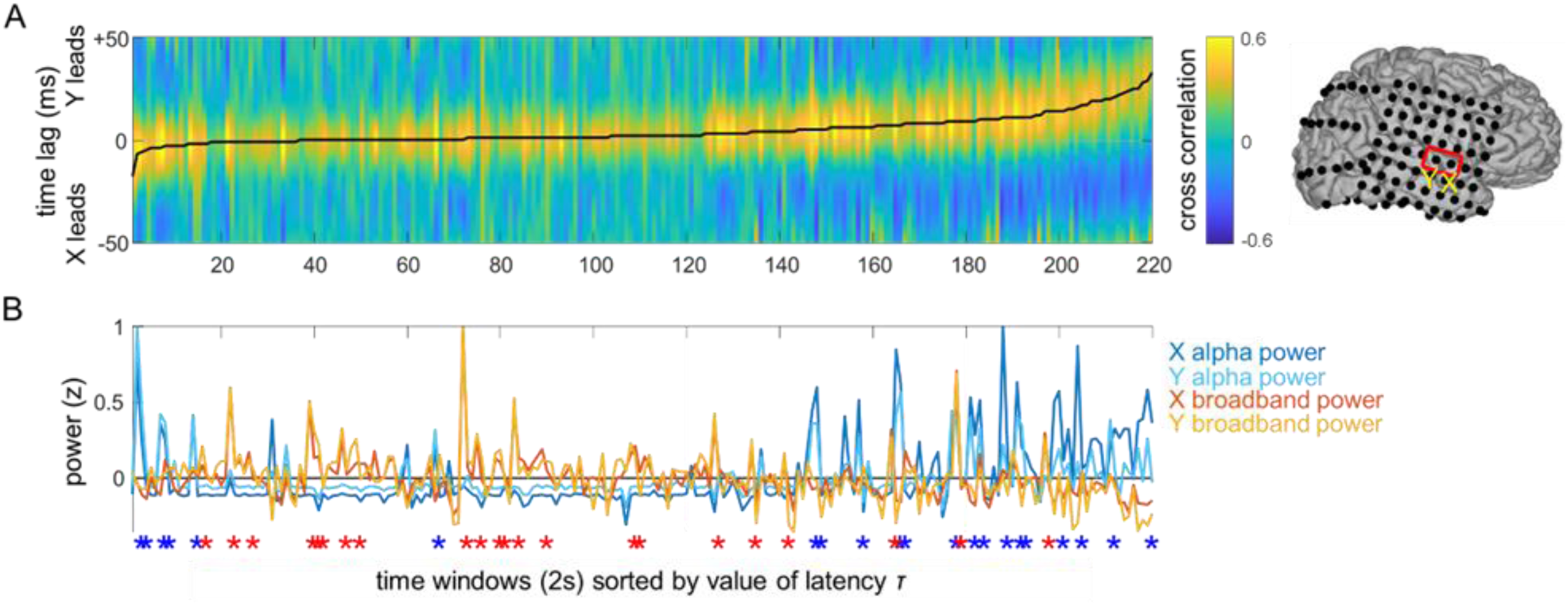
Latency and power changes for an electrode pair in the middle STG. Time windows are sorted by value of Latency τ between electrodes *X* and *Y*. For each channel pair, we arranged the time windows in the increasing order of the latency *τ*, and measured mean alpha-band (10 Hz) power and mean broadband (65+ Hz) power across channels for each corresponding time window. **(*A*)** Cross correlation between two chosen channels is shown for each time window, sorted by the value of latency *τ*. Black line denotes the inter-electrode latency *τ*. **(*B*)** Alpha-band (7-14Hz) and broadband power in each channel for each 2s time window. Blue asterisks denote alpha peaks and red asterisks denote broadband power peaks (top 10% of values). The latency *τ* is correlated with alpha power and anti-correlated with broadband power.

The relationship between inter-channel latency and low-frequency power was observed across the parietal, temporal and peri-Rolandic cortex (Fig. 3*A*). Because latencies can only be reliably estimated for regions which exhibit coupling in their dynamics, we focused on “stable channel pairs”, which were defined as sites within 25 mm of one another which exhibited peak Pearson cross correlations > 0.3 for 80% of the time windows. Using the stable channel pairs, we then estimated the “latency flow” across the cortical surface by spatially and temporally averaging the vectors of flow between stable pairs (see Methods). Comparing the latency flow for the time windows with the top 10% and bottom 10% of global alpha power (alpha power averaged across all channels), it is apparent that inter-channel latencies and inter-channel correlations increased with alpha power (Fig. 3*A*, left, longer latencies indicated by larger arrows, stronger correlations indicated by warmer colors). Similar results are observed for all low-frequency bands (Supplementary Figs. S1-S3, Table S2). This latency effect was confirmed for each of the 10 participants individually (t-test comparing latencies for top 10% power and bottom 10% power, all p’s < 0.05 individually), while the correlation strength effect was present in 8 of the participants. We observed a complimentary (and opposite) effect for high-frequency broadband power (65 + Hz): the inter-channel latencies decreased as global broadband power increased (Supplementary Fig. S4, Supplementary Table S2).

**Fig. 3.**
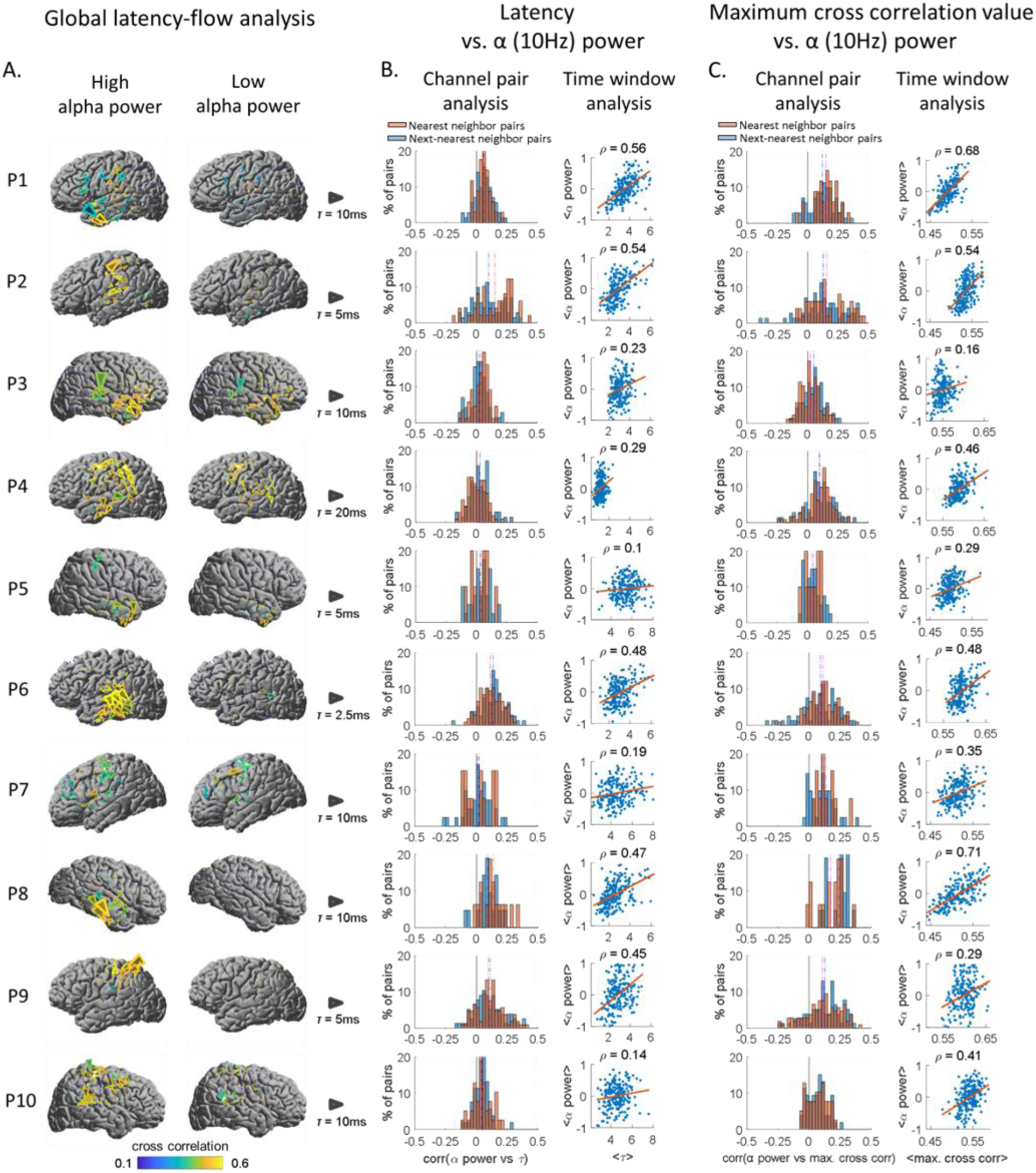
Latencies and coupling strengths increase with low-frequency power. **(*A*)** *Global latency-flow analysis.* Latency-flow patterns for high alpha power time windows and low alpha power time windows are shown. Top 10% of the windows yielding highest alpha power and bottom 10% of the windows yielding lowest alpha power are chosen, and latency flows are computed across the chosen windows respectively. Yellower color of arrows denotes higher cross correlation. Larger size of arrows denotes longer latencies. The size of arrows on each flow map is scaled for readability: the grey arrow next to each participant’s brain map denotes scale of latency flow arrows. **(*B*)** *Time delay vs. alpha power.* Channel pair analysis: Correlation between latencies of pairs and global alpha power for each pair across time windows are computed and the distribution is shown in histograms. Red histogram represents nearest neighbor pairs and blue histogram represents next-nearest neighbor pairs. The distribution is positively skewed. Time window analysis: Mean latencies across pairs and mean global alpha power for each time window are computed and shown as scatterplots. The distributions yield positive correlation. **(*C*)** *Maximum cross correlation vs alpha power.* Channel pair analysis: Correlation between maximal cross correlation of pairs and global alpha power for each pair across time windows are computed and the distribution is shown in histograms. The distribution is positively skewed. Time window analysis: Mean maximal cross correlations across pairs and mean global alpha power for each time window are computed and shown as scatterplots. The distributions yield positive correlation.

Consistent with prior reports ^18,19^, we observed that that most flow arrows pointed anterior-inferior in the temporal and parietal lobes, indicating that parietal and posterior temporal dynamics were time-advanced relative to anterior and inferior temporal sites (Supplementary Fig. S5-S6). Of the 8 participants whose coverage included the temporal lobe, 7 exhibited mean flow angles < π/2 radians relative to the posterior-anterior axis along the Sylvian Fissure.

Having found that global flow patterns differed between the smallest and largest oscillation magnitudes (Fig. 3*A*), we next tested the latency changes at the level of individual pairs of electrodes across all values of oscillation magnitude. For each channel pair, we computed the Spearman correlation between the latency and the global LF power across time windows. We aggregated these values across channel pairs to generate a distribution of correlations. For every participant, the distribution had a positive mean (Fig. 3*B*, Channel Pair Analysis, mean of subject-wise mean Spearman ρ = 0.10), indicating a consistent positive correlation between channel pair latency and the global LF power. This phenomenon was observed for nearest neighbor channel pairs (distance < 12.5 mm, Fig. 3*B*, red) and for next nearest neighbor channel pairs (12.5mmm < distance 25 mm, Fig. 3*B*, blue). Similar results were obtained when we estimated oscillation magnitude using the local LF power nearby each pair of channels, rather than the global LF power averaged across channels (see Methods). Finally, taking a more spatially averaged view, we observed that the mean latency across all electrodes increased within time windows with greater global LF power (Fig. 3*B*, mean latency analysis, mean Spearman ρ = 0.37).

These measurements indicate that the relationship between latency and oscillation amplitude was robust and consistent across measurement methods. Note that it was not possible to test this relationship within electrodes whose couplings were transient or weak (inter-electrode r < 0.3, making up 42% of nearest neighbor channel pairs and 83% of next-nearest neighbor channel pairs), because it was not possible to reliably estimate the inter-electrode delays in such electrodes.

### Inter-channel correlations increase with low-frequency oscillatory amplitude

In parallel with the latency effects described above, we also observed that higher amplitude LF power was associated with larger inter-electrodes correlations. This effect is qualitatively apparent in Fig. 3*A*, where greater correlation magnitude (warmer arrow colors) were observed for the higher levels of LF power (left column). Quantitatively, the histogram of correlations between LF-power and inter-channel correlation were positively shifted for all participants (Fig. 3*C*, channel pair analysis, mean Spearman ρ = 0.20 when correlating LF power and coupling magnitude) and the spatial mean of the inter-channel correlation also increased with the LF power (Fig. 3*C*, mean Spearman ρ = 0.45). Once again, the effects associated with LF power were very similar when measured with globally averaged LF power and with the LF power specific to a channel pair (Supplementary Fig. S7). Moreover, these effects also persisted when we included next-nearest neighbor electrode pairs (Fig. 3*B*, *C*).

### Stronger oscillations are associated with fewer zero-lag couplings

Did the changes in latency also corresponded to a shift away from isochronous (zero-lag) synchronization? On the one hand, LF power changes might covary with a shift in non-zero delays (e.g. a shift from 5 to 10 ms latency), but on the other hand, LF power changes may covary with a shift from an isochronous synchrony to a lagged synchrony (e.g. from 0ms to 5 ms latency), with distinct functional implications ^17^. Defining zero-lag as any inter-regional coupling with a latency estimated in the range [-2, 2] msec, we found (in all participants individually) that increases in LF power were associated with decreases in the number of zero-lag pairs (Supplementary Fig. S8). Thus, as LF power increased, the absolute number of zero-lag couplings decreased, even while the number of coupled pairs increased.

### Latency fluctuations are weakly locked to the stimulus in the auditory pathways

Given that fluctuations in cortical oscillations can be reliably locked to an auditory narrative stimulus ^24,25^, can the inter-regional latency patterns also be locked to an auditory narrative? To answer this question, we first identified pairs of channels which both exhibited a consistent LF power time course across repeats of the narrative stimulus. For example, the channels highlighted in Fig. 2*A* exhibited reliable single-trial LF fluctuations across repeats. For these channels, we defined the *latency reliability* as the correlation between latency values in corresponding 2s windows for run 1 and run 2 of the auditory stimulus (Fig. 4*B*, *C*). For this pair, we observed that *latency reliability* (Spearman ρ = 0.24 across run 1 and 2), was lower than *LF power reliability* (defined as the Spearman correlation between LF power values in corresponding 2s windows for run 1 and run 2) in channel X (Spearman ρ = 0.66 across run 1 and 2) or channel Y (ρ = 0.45).

**Fig. 4.**
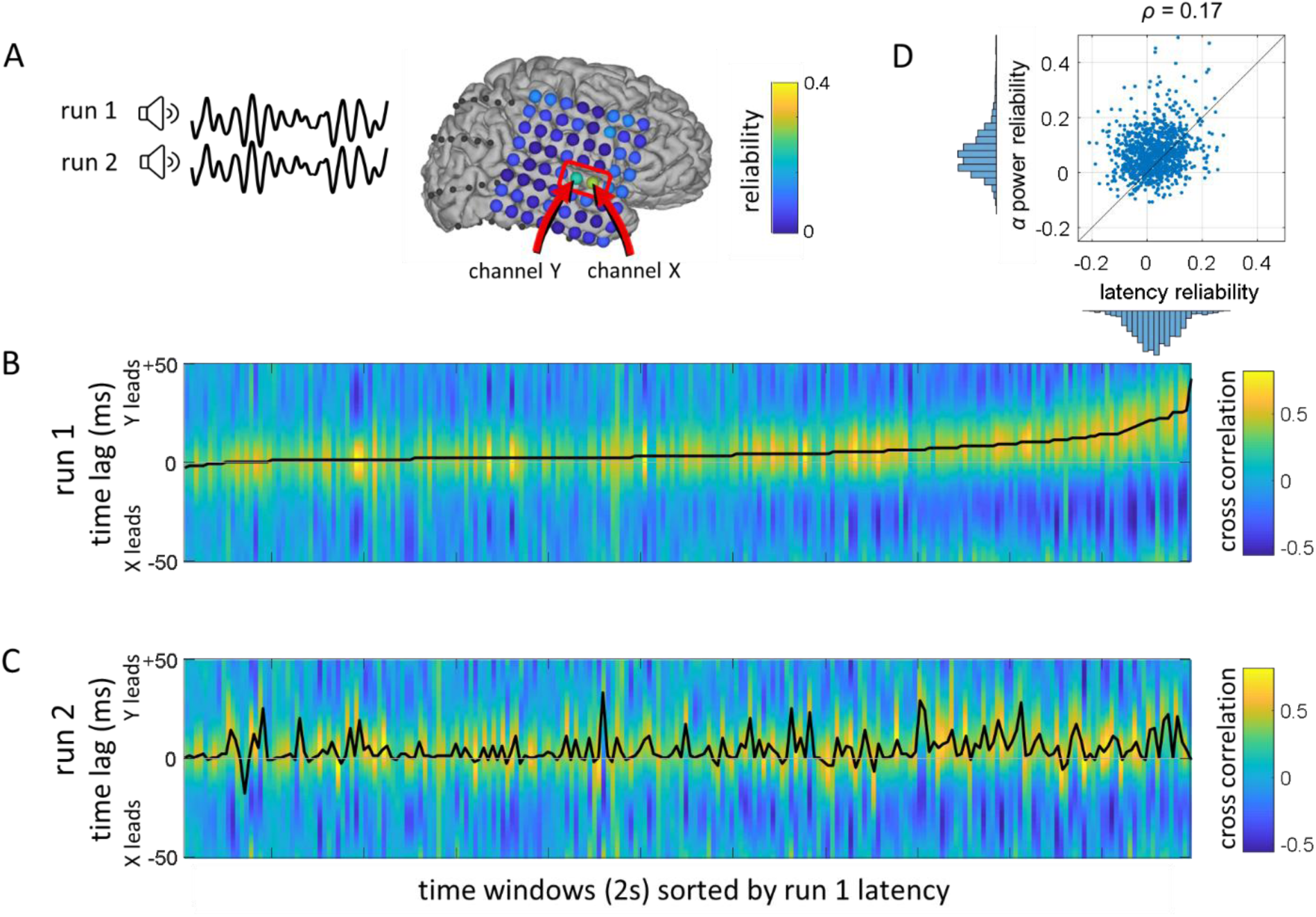
Reliability of latency patterns between run 1 and run 2. **(*A*)** Two runs of the same auditory stimulus were applied to a participant., and channels with high reliability between two runs were chosen. **(*B*)** The latency pattern for run 1 is shown in the order of increasing latency. **(*C*)** The latency pattern for run 2 is shown in the same order used in run 1. The latency reliability between two runs was 0.25 for the chosen channel pair. **(*D*)** Latency reliability vs. LF power reliability for all channel pairs from 10 participants.

Latency reliability was low even in sites entrained to the auditory stimulus, suggesting that fluctuations in latency were more strongly controlled by endogenous processes than by our auditory narrative. For each channel pair with measurable latencies, we compared the latency reliability of that pair against its *LF power reliability* (Spearman correlation between LF power time courses for run 1 and run 2) and its *broadband power reliability* (defined similarly). Across all 10 participants, the latency reliability and LF power reliability were only weakly related (Fig. 4*D*, Spearman ρ = 0.17; individual participant LF and broadband data in Supplementary Figs. S9-S10). We found that 12% of the pairs yielded LF power reliability greater than 0.2, and 25% exhibited broadband power reliability greater than 0.2, but less than 5% of the pairs yielded the corresponding levels of reliability in their latency time courses. The pairs exhibiting both reliability in LF time courses and in latency patterns were mostly located in in early auditory pathways of the superior temporal cortex.

### A coupled oscillator model identifies inter-node influence as key parameter

To probe possible mechanisms underlying the relationships between latencies and global oscillation patterns, we constructed a cortical network model. The model was composed of 78 nodes linked according to inter-regional connections in the human brain ^26^ (Fig. 5*A*). We employed Stuart-Landau oscillators to represent the neural mass activity at each node, because these oscillators follow the normal form of the Hopf bifurcation, providing the simplest model that captures the amplitude and phase dynamics of neural systems near a bifurcation point ^27,28^. The dynamics of each node in the model arise from a combination by (i) intrinsic phase and amplitude dynamics determined by the natural frequency and amplitude assigned to each node and (ii) inter-node influences from anatomically connected neighbors. We fixed the parameters of the intrinsic dynamics, drawing the natural frequencies for each node from a Gaussian distribution with mean of 10 Hz and s.d. of 1 Hz. We varied the coupling strength, S, which globally determines the magnitude of each node’s influence on its anatomical neighbors, between S = 0 (no coupling) and S > 1 (very strong coupling).

**Fig. 5.**
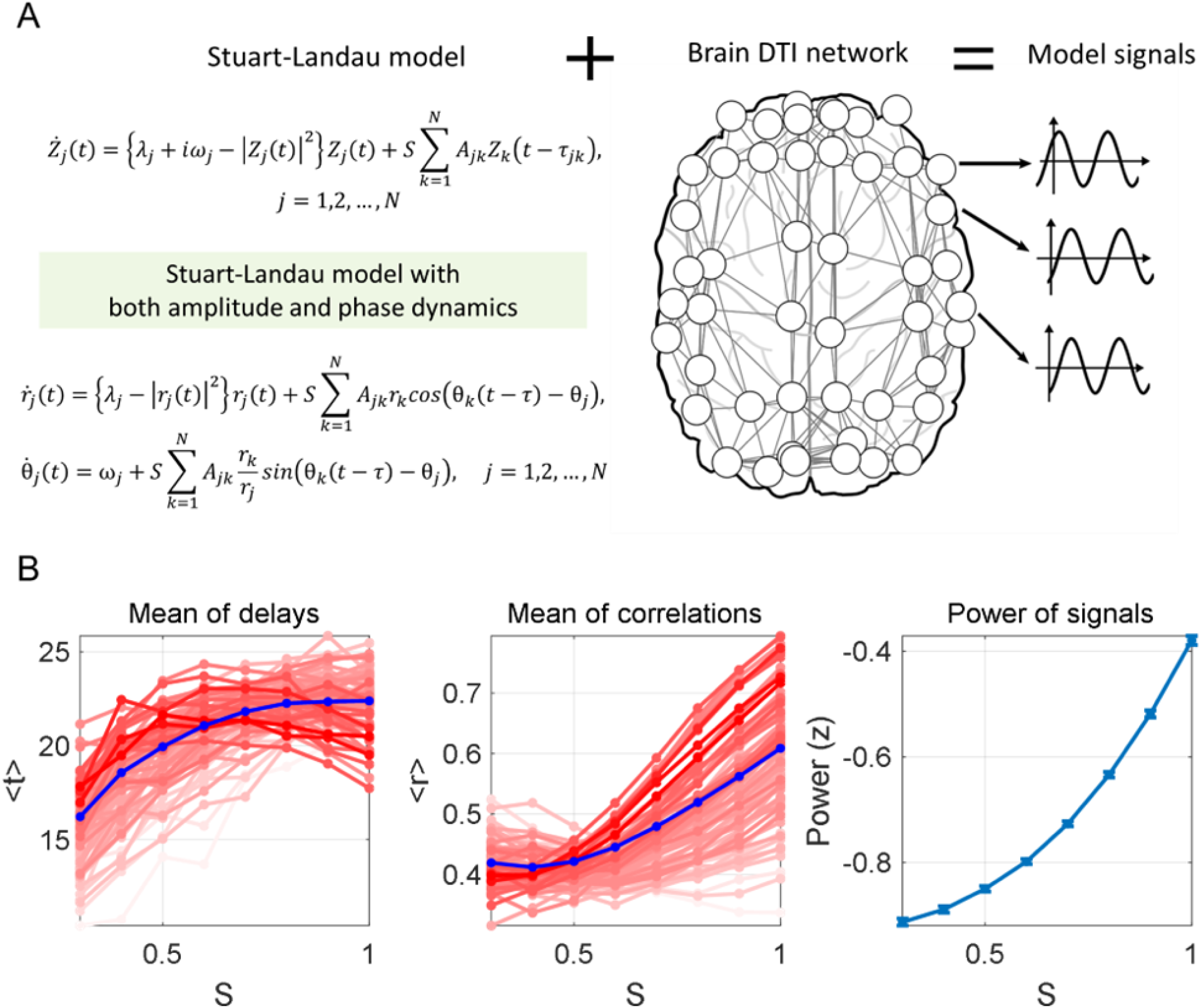
Coupled oscillator model on brain network. **(*A*)** Stuart-Landau coupled oscillator model is applied to diffusion tensor imaging (DTI) structural brain network consisting of 78 nodes. **(*B*)** Mean delay across the nodes, mean of maximal cross correlations across the nodes, mean of power of the nodes are shown. Below a certain critical coupling strength S (∼1.0), all three measures are positively correlated with each other as the coupling strength S increases.

For intermediate values of the inter-node influence, 0.4<S<1, we observed a match to the empirical phenomenon: there was a positive correlation between the mean latency across nodes, the mean peak cross correlation, and the mean oscillatory amplitudes (Fig. 5*B*). When the inter-node influence (modelled via the coupling strength S) was near zero, the nodes exhibited a uniform distribution of relative phases, because their dynamics were independent. Thus, for S near zero, there was approximately zero correlation between the dynamics of neighboring nodes over many cycles. For strong coupling strengths (S > 1) the nodes became tightly locked with each other over entire cycles and pulled one another into zero-lag synchrony. Thus, strong coupling was associated with decreased latency and increased correlation between network neighbors. However, in the intermediate range of coupling values (0.4<S<1), the oscillators became phase-locked for only a portion of each oscillation cycle, and with non-zero phase difference. In this range, the mean inter-node latency increased with S, as larger values of S within this range were associated with a larger proportion of time spent phase-locked within each cycle.

What other classes of models might explain the experimentally observed correlation between latency and low-frequency power? Moving towards more abstracted models, with less biological connection, the phenomenon can be explained by models that posit a change in the relative magnitude of delayed and simultaneous inter-node couplings (details in Supplementary Text). For example, we considered a generic model composed of two time series processes, each time series expressed as the sum of two processes: a common process (coupled with a fixed zero delay) and an oscillatory process (coupled across sites with a fixed nonzero delay). As we increase the amplitude of the coupled oscillatory process, the observed inter-regional latency and the oscillatory power will increase in tandem (Supplementary Text, Supplementary Fig. S11). This simplified model can be understood as an abstraction of the Stuart-Landau model processes (Fig. 5). However, regardless of whether the latency shifts arise from the Stuart-Landau mechanisms (an increase in inter-regional global coupling) or via this more abstracted model, the neurobiological outcome does not change: moments of peak excitability shift from zero-lag to nonzero-lag as low-frequency oscillations increase.

## Discussion

Recording intracranially from the human cerebral cortex, we observed that increases in low-frequency power (4-14Hz, especially 6-10Hz) were associated with stronger and more delayed couplings between regions. In time windows when high-frequency broadband power (65+ Hz) was increased we observed the opposite effect: weaker inter-regional couplings and a larger proportion of zero-lag coupling. These changing coupling patterns were observed across the lateral cerebral cortex and were associated with both locally and globally-measured oscillatory processes. Moreover, the increases and decreases in coupling latency were mostly endogenous: when we repeated the same auditory speech stimulus, only a small number of auditory-locked sites exhibited reliable latency patterns across repetitions. Using computational models composed of amplitude-and-phase-coupled oscillators, we found that the empirical changes in coupling magnitude and latency could be explained by increases and decreases in the effective cortico-cortical influence.

What are the functional consequences of the continual waxing and waning of low-frequency oscillatory processes in the human brain? When oscillations are stronger, this is often associated with a decrease in perceptual sensitivity and an increase in the relative strength of long-range communication^29^ and perhaps top-down signaling in particular^30,31^. Indeed, low-frequency oscillations have been linked to an idling state and “maintaining the status quo”^32,33^. Here, we show that not only are stronger oscillations associated with stronger coupling between brain regions, but we also find that stronger oscillations are associated with longer delays between regions. Thus, when oscillations are weaker, the couplings are weaker and synchronous; when oscillations are stronger, the couplings are stronger and delayed. Indeed, when the oscillations are especially strong, then many nearby sites may exhibit a shift to a lagged coupling state, and this may manifest in the form of traveling waves across the cortical surface ^19,34,35^.

Our amplitude-and-phase based model can be understood as a generalization of the phase-based model proposed by Zhang et al., 2018 to account for cortical traveling waves. Zhang et al. (2018) showed how traveling waves could be generated by gradients of intrinsic frequencies across a lattice-like cortical sheet. Our model can produce the same traveling wave effects, but it leads to two further predictions. First, our model predicts that continuous changes in the amplitude of the oscillations are related to continuous changes in latency, even in the absence of gradients of intrinsic frequency. Secondly, our model indicates that a node’s degree (number of connections) will influence whether it leads or lags its neighbors (Woo et al., 2021).

What are the functional implications of coordinated shifts between zero-lag and delayed couplings? The topographies of flow were somewhat variable across participants, consistent with the reports of Zhang et al. (2018), but there were some reliable patterns. In particular, when LF oscillations were strong within the temporal-parietal auditory pathways, the field potentials in higher stages of processing (parietal and posterior temporal cortex) were time-advanced relative to more peripheral regions of the auditory pathway (middle superior temporal cortex) (Fig. 3 and Supplementary Fig. S1-S2). If we assume that there is a “preferred potential” at which the excitability of cortico-cortical projections is increased ^15,36^, and that inter-regional axonal conduction delays are 5-10ms ^37^, then the oscillations will modulate the effectiveness of signaling between regions. For example, when 10 Hz oscillations are elevated, leading to a 5-10 ms time-lead (Fig. 3), then the peak excitability phase of higher order regions will generate spikes that arrive at the peak excitability phase of the low-order regions. Conversely, in the same state, spikes arising from the peak excitability phase of lower-order regions will arrive after the peak excitability phase of the higher order regions. Thus, increases in low-frequency oscillations could increase the relative strength of top-down signaling in the human brain, consistent with theoretical and empirical recordings from non-human primates ^30,38^.

We hypothesize that the changes in cortico-cortical delays observed here, in conjunction with changes in oscillatory power, are regulated by spatially nonspecific ascending projections. These signals could arise from controllers of physiological arousal or from nonspecific thalamocortical projections. First, a nonspecific regulation is consistent with the observation that the latency between cortical regions was not only increased in relation to the alpha power in those regions, but also with non-local alpha power measured centimeters away (Fig. 3). Second, we observed only a weak association between the latency patterns and properties of the auditory stimulus that was presented. Third, arousal changes are known to regulate the same oscillatory processes that we measured here ^20,21^. Fourth, multi-second latency changes coupled to physiological arousal have been reported in humans using fMRI ^23^, and latency changes in cats have been observed in relation to spontaneous changes in their attentive state, attributed to ascending modulatory projections ^12^. Finally, our mathematical model was able to account for the changes in latency via a global (nonspecific) modulation of coupling strength.

Altogether, the data and models suggest that human cortical dynamics reliably transition between near-zero latency states (associated with suppression of alpha-band power and stronger broadband high-frequency power) and longer latency states (associated with increases of alpha-band power and decreases in high-frequency broadband power). The latency changes were observed in widespread temporal, parietal and somatomotor sites; thus, these delay-fluctuations may be associated with distinct large-scale functional states. Although oscillations have been proposed to have multiple roles in organizing cortical communication, the present findings emphasize the importance of slow fluctuations in the amplitude of these oscillations, which appear to mediate transitions between states of latency and dynamical flow across the surface of the human brain. These slow fluctuations may correspond to the large-scale neural state changes observed using fMRI ^39,40^.

Two primary limitations of this work are, firstly, that we have not yet mapped the latency fluctuations across a wider range of cognitive and perceptual tasks; and secondly, that we have access to only a subset of brain regions from a clinical population. In relation to the range of stimuli and task states: a natural next step would be to measure these transitions in a sustained attention or vigilance task ^4,41–43^, or in the kinds of state-transitions tasks which are associated with gradual large-scale network transitions in neuroimaging ^44–46^. In relation to the fact that these recordings are from epilepsy patients with coverage primarily in temporal, parietal and peri-Rolandic cortices: these findings may be detectable within a wider range of areas in a neurotypical population using sensor-space analysis of high-density EEG. At the same time, given the earlier report of latency state transitions in the cat brain ^12^, it may be equally fruitful to search for latency state-transition in rodent or nonhuman primate brains, where high-resolution recording technologies will make it possible to identify the connection to modulations of the underlying firing rates ^47^.

## Methods

### Participants

10 patients (7 female; 19-50 years old) were recruited from the Surgical Epilepsy Program at Toronto Western Hospital (Ontario, Canada) from the pool of all patients evaluated for neurosurgical treatment of medically refractory epilepsy. The clinical and demographic information of participants is summarized in Table S1.

### Data acquisition

Signals were recorded from 64 contact grids. Electrode placement was solely based on clinical criteria. For data acquisition patient connectors were transferred to a separate research amplifier system (NeuroScan SynAmps2; Compumedics, Charlotte, NC). Data was recorded at 5 kHz and 0.05 Hz hardware band-pass filtering. Event markers from the stimulus presentation were sent to the SynAmps2 through a parallel port. The SynAmps2 also recorded the participant’s overt spoken response and the surrounding acoustics through a microphone connected to a Tube MP Studio V3 preamp (ART ProAudio, Niagara Falls, NY). Electrodes were localized with the freely available iELVIS toolbox for MATLAB (http://ielvis.pbworks.com).

### Data preprocessing

Electrodes exhibiting artifacts, epileptiform activity, excessive noise or no signal were excluded from the analysis by offline visual inspection. As the raw voltage signals of some recording sessions in a given patient exhibited different means and variance, preprocessing was done for each session separately (about 7 minutes and 20 seconds of recording). Raw voltage signals were down-sampled to 1000 Hz. To minimize filter artifacts, all filtering was performed on continuous data. Data was high-pass filtered at 0.1 Hz (Butterworth, order 4) and low-pass filtered at 200 Hz (Butterworth, order 4). Line noise was removed with a bandstop filter (Butterworth) from 59.5-60.5 Hz, 119.5-120.5 Hz and 179.5-180.5. Similar to an average-rereferencing approach, the mean voltage signal across all remaining channels of a given subject was projected out from each individual channel by using linear regression. The voltage trace of each channel was z-scored, including only the trial periods for calculating the mean and standard deviation. To avoid edge artifacts, filtering was always performed on longer segments of continuous data.

### Computation of power time courses

We estimated time courses of signal power using Morlet wavelets. For the frequency range 65-200 Hz, power time courses were computed separately for each frequency in steps of 5 Hz. 120 and 180 Hz were excluded as harmonics of line noise. We then took the logarithm of each power time course estimate.

These estimates were then z-scored. Finally, we computed the mean across each of the frequency-specific z-scored time courses to yield a single “broadband” 70-200 Hz power time course. Because of the long duration of our trials, we observed transient motor-related artifacts due to swallowing that contaminated all channels in the same frequency range as broadband power. We removed this artifact in two steps. First, we generated a temporally sparse noise regressor by identifying deviants from the median broadband power time course across all channels in a recording session. At times when the median broadband power across channels was less than three times the interquartile range from its median value, the noise regressor was set to zero; at all other times (i.e., during bursts), the noise regressor was set equal to the mean broadband power across channels. We then projected out this noise regressor from the continuous broadband power time course by taking the residual from a linear regression of the noise regressor on the broadband power signal in each channel. Finally, the broadband power time courses were smoothed with a 500 ms Hamming window.

For lower frequency power bands, the peak was chosen as 2 Hz for delta power band, 5 Hz for theta power band, 10 Hz alpha power band, and 20 Hz for beta power band. Using Morlet wavelets, power time courses were computed for each frequency using a wave-width of 6. The same noise-removing processes were applied as used for the broadband power computation. Again, each power time courses were smoothed with a 500 ms hamming window.

### Cross correlation analysis

Each participant listened to two repetitions of a 7-minute auditory stimulus with a narrative. First, in sliding 2 second windows, we computed the cross-correlation of the raw voltage signal between adjacent sites (channels within the distance of 25 mm from each other). For each channel-pair in each time window, we identified the time lag of maximal inter-electrode correlation and defined it as the latency τ between the channels. For each channel and 2 second window we compared the cross-correlation with the mean alpha (7-14 Hz) power and mean broadband (65-200 Hz) power of the channels in the pair.

### Network analysis

To test whether the latency-alpha power relationship is consistent across channel pairs, we observed global latency pattern changes between high alpha power time windows and low alpha power time windows. First, we chose top 10 % of time windows with highest alpha power (20 windows). For each channel and each time window, we applied vector-summation of the latencies from the channel to all the other channels within the threshold distance (25mm) to derive a “latency-flow” of the channel for the corresponding time window. Then we applied again vector-summation of each latency-flow across the chosen time windows to finally derive time-averaged latency-flow for each channel. We also computed the time averaged cross correlation value for each channel in a similar way. For each channel and each time window, we computed spatially averaged maximal cross correlation between the channel and all the other channels within the threshold distance (25mm) to derive a spatially averaged maximal cross correlation of the channel for the corresponding time window. Then we again averaged across the chosen time windows to derive time-averaged maximal cross correlation for each channel.

After we derived these latency-flow and averaged maximal cross correlations for the top 10% alpha power time windows, we repeated the same process for the bottom 10% of time windows with lowest alpha power (20 windows).

### Models

We develop a couple oscillator model with both phase and amplitude dynamics, in the form of Stuart-Landau equation (Equation (1.1) and (1.2)).

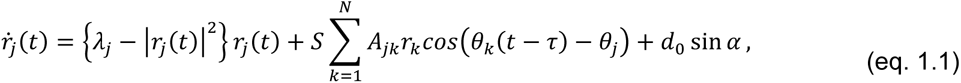

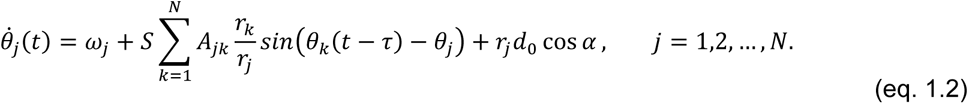

Here, Equation (1.1) represents amplitude dynamics and (1.2) represents phase dynamics. *r_j_(t)* is the amplitude and *θ_j_(t)* is the phase of an oscillator *j* at time *t*. *S* is a parameter controlling the global coupling strength in the system, and *A_jk_* denotes the coupling strength between oscillator *j* and *k*. *ω_j_* is the intrinsic frequency of the *j* and *λ_j_* is the bifurcation parameter controlling how fast the trajectory converges to stability; in this model we consider *λ_j_ = λ > 0* for all *j = 1,2,…, N,* and in such case 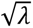 is considered as the “intrinsic amplitude” to which the oscillator converges in the absence of the coupling. *d_0_* sinα and *d_0_* cos *α* represent the effect of constant “external field” applied to all the oscillators. Every oscillator with Hopf bifurcation reduces into Stuart-Landau model in the vicinity of the bifurcation, and neural oscillators are known to exhibit such bifurcations. The Equations (1.1) and (1.2) are generalizations of Stuart-Landau model in a sense that they reduce to Stuart-Landau model without the coupling terms.

## Data availability

The patient data were collected under a Canadian clinical research protocol (Research Ethics Board, University Health Network) which does not allow deposition in public data archives. Data and scripts can be shared upon request.

## Acknowledgments

The authors gratefully acknowledge the support of the National Institutes of Mental Health (grant R01MH119099 to C.J.H.; grant R01 MH111439 to C.E.S. with subaward to C.J.H.), Institute for Basic Science of Korea (Young Scientist Fellowship IBS-R015-Y3 to J.-Y.M.), and the Alfred P. Sloan Foundation (research fellowship to C.J.H.) as well as the Toronto General and Toronto Western Hospital Foundation. The authors would also like to thank Victoria Barkley for assistance with data collection; David Groppe for support with the iElvis toolbox; and to the staff of the epilepsy monitoring unit at Toronto Western Hospital for logistical support. Finally, our great thanks to the patients who participated in this study.

## Author Contributions

J.-Y.M. and C.J.H. designed research; J.-Y.M. and C.J.H. performed research; J.-Y.M., K.M., and C.J.H. analyzed data; J.-Y.M. performed modeling; J.-Y.M., K.M., C.E.S., and C.J.H. wrote the paper; and T.A.V. supported the data acquisition.

## Competing Interest Statement

The authors declare no competing interest.

## Supplementary Information

This paper accompanies Supplementary Text, Supplementary Figures S1-S11, and Supplementary Tables S1-S2.

## Notes

### Competing Interest Statement

The authors have declared no competing interest.

### Summary of Updates

Bibliography updated

