## Supplementary Material for "Inter-regional delays fluctuate in the human cerebral cortex"

#### Abstracted Models with Varying Signal-to-Noise of Oscillatory Processes

##### Model 1: Delayed oscillatory coupling

Can the change in delay be explained by changes in the signal-to-noise ratio of an oscillatory process that is associated with a fixed time delay? In particular, suppose that the signal measured in each channel is expressed as a sum of an oscillatory signal component and a random noise process (denoted as $R_1$ and $R_2$, respectively):

$$X(t) = A\sin(\omega\, t) + \; B\, R_1(t),$$
$$Y(t) = C\sin\big(\omega(t-\tau^*)\big) + D\, R_2(t). \qquad \text{(eq. 2.1)}$$

As the parameters *A* and *C* increase relative to *B* and *D*, the magnitude of the measured oscillation will become larger, and the empirically estimated $\tau$ value will converge toward the true coupling delay, $\tau^*$.

However, we confirmed using simulations that, under such a model, when the oscillatory signal amplitudes are very low, the cross-correlograms are uniform random numbers on the range [$-\tau_{max}$, $\tau_{max}$], where $\tau_{max}$ is the largest delay measured. For $\tau_{max}$ = 50ms, as used in our study, this would lead to a mean absolute delay of 25 milliseconds. In contrast, we empirically observed tau values near zero when the oscillatory amplitude was small.

##### Model 2: Distinct oscillatory couplings at nonzero lag and at zero lag

A second possible mechanism by which the lags could vary with signal-to-noise ratio is if following two processes are combined: First, an oscillatory process expressed with a fixed time lag across both channels (as in Model 1 above), and second, a common drive process that is expressed with zero lag across both channels. Such a model can be expressed as:

$$X(t) = A\sin(\omega\, t) + B\,\sin(t) + \; C\, R_1(t),$$

$$Y(t) = A\sin\big(\omega(t-\tau^*)\big) + B\,\sin(t) + \; C\, R_2(t). \qquad \text{(eq. 2.2)}$$

We can consider a case where we increase parameter *A* relative to *B* and *C* (where *B* is sufficiently larger than *C*). In such case, the measured value of $\tau$ increases from zero to the original coupling delay $\tau^*$. By addition of trigonometric function, it can be shown that when *A* = *B*, the delay $\tau$ equals $\tau^*/2$:

$$X(t) = 2A\sin(\omega t) + \; C\, R_1(t),$$
$$Y(t) = A\cos\left(\tfrac{1}{2}\omega\tau^*\right)\sin\left(\omega(t - \tfrac{1}{2}\tau^*)\right) + \; C\, R_2(t). \qquad \text{(eq. 2.3)}$$

As *A* increases further (*A*>*B*), $\tau$ increases further (*SI Appendix* Fig. S11). This case can be considered as a special case of our model in the main text. We can consider our coupling term in our model as the time delayed term in eq. 2.2, where in our model the time delay also varies depending on the coupling strength.

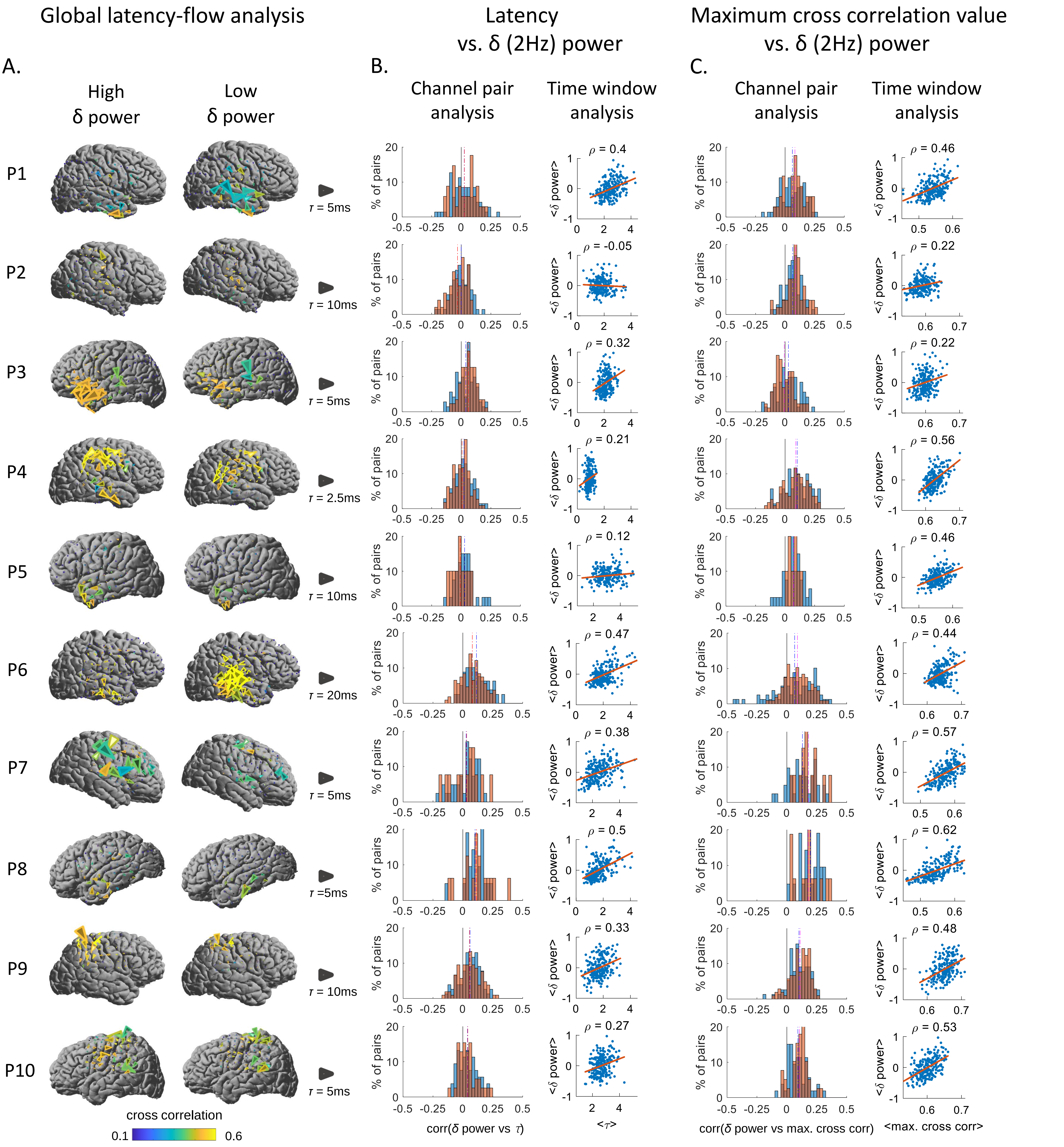


**Fig. S1. Latencies and coupling strengths increase with delta band power. (*A*) *Global latency-flow analysis*.** Brain surfaces illustrate latency flow maps averaged across time windows with high values (top 10% ) and low values (bottom 10%) of global delta-band power. Top 10% of the windows yielding highest delta power and bottom 10% of the windows yielding lowest delta power are chosen, and latency flows are computed across the chosen windows respectively. Yellower arrows denotes higher cross correlation. Larger size of arrows denotes longer latencies. The size of arrows on each flow map is scaled for readability: the grey arrow next to each participant's brain map indicates scale of latency flow arrows for that participant. (***B***) ***Time delay vs. delta power***. *Channel pair analysis*: For each pair of channels, the inter-channel latency and global delta power are correlated across time windows and the distribution of correlation values is shown in histograms. Red histograms represent nearest neighbor channel pairs and blue histograms represent next-nearest neighbor pairs. The distribution is positively skewed. *Time window analysis*: The mean latencies across pairs and mean global delta power for each time window are computed and shown as a scatterplot, with one point per time window. (***C***) ***Maximum cross correlation vs. delta power.*** *Channel pair analysis*: For each pair of channels, the inter-channel peak cross-correlation value is correlated across time windows with  global delta power. The resulting correlation values are shown in the histogram. *Time window analysis*: For each time window, the spatial mean of the inter-channel correlations is computed along with the global delta-band power and shown as a scatterplot, with one point per time window.. The distributions yield positive correlation.

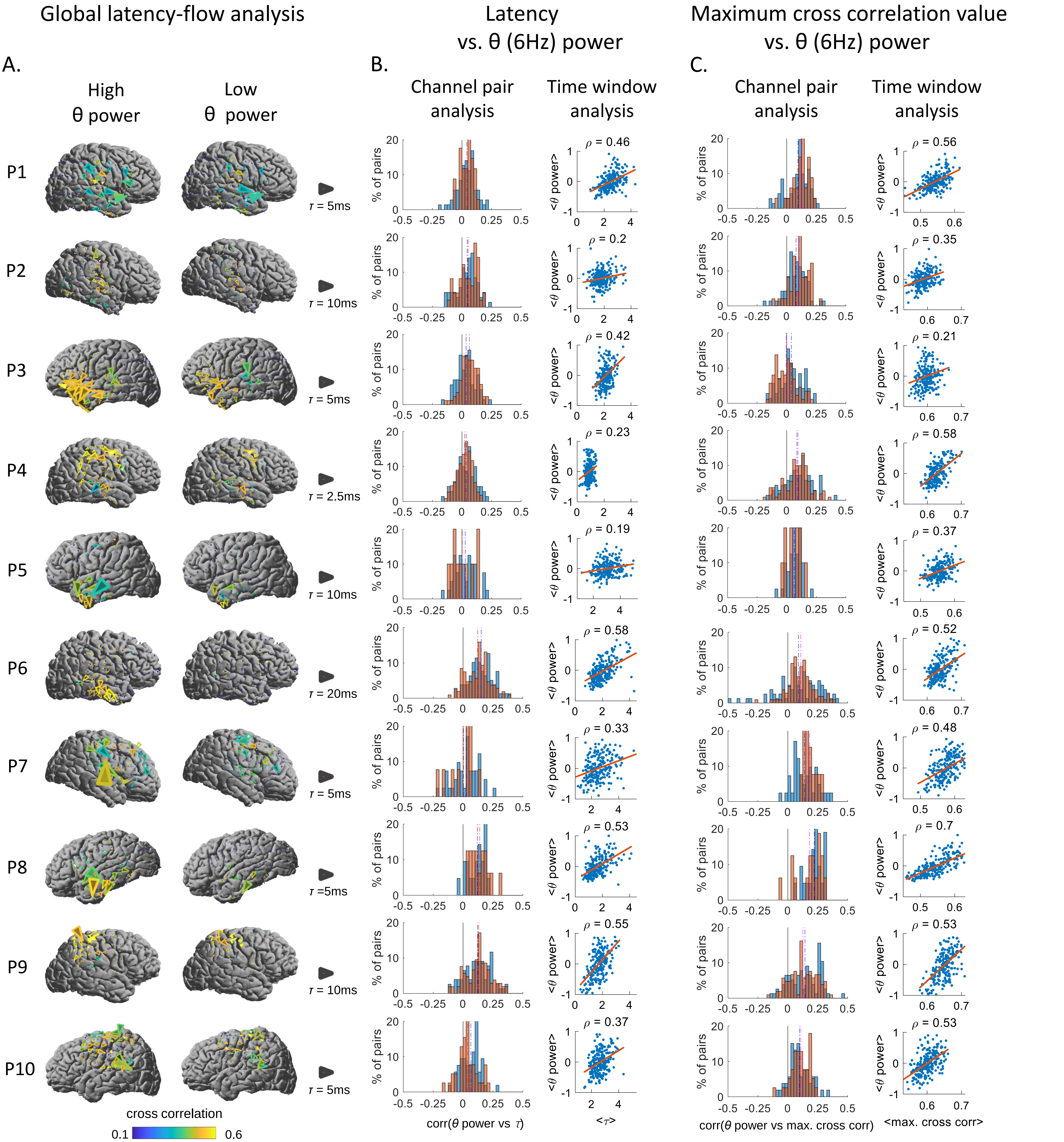


**Fig. S2. Latencies and coupling strengths increase with theta band power. (*A*) *Global latency-flow analysis*.** Brain surfaces illustrate latency flow maps averaged across time windows with high values (top 10% ) and low values (bottom 10%) of global theta-band power. Top 10% of the windows yielding highest theta power and bottom 10% of the windows yielding lowest theta power are chosen, and latency flows are computed across the chosen windows respectively. Yellower arrows denotes higher cross correlation. Larger size of arrows denotes longer latencies. The size of arrows on each flow map is scaled for readability: the grey arrow next to each participant's brain map indicates scale of latency flow arrows for that participant. (***B***) ***Time delay vs. theta power***. *Channel pair analysis*: For each pair of channels, the inter-channel latency and global theta power are correlated across time windows and the distribution of correlation values is shown in histograms. Red histograms represent nearest neighbor channel pairs and blue histograms represent next-nearest neighbor pairs. The distribution is positively skewed. *Time window analysis*: The mean latencies across pairs and mean global theta power for each time window are computed and shown as a scatterplot, with one point per time window. (***C***) ***Maximum cross correlation vs. theta power.*** *Channel pair analysis*: For each pair of channels, the inter-channel peak cross-correlation value is correlated across time windows with global theta power. The resulting correlation values are shown in the histogram. *Time window analysis*: For each time window, the spatial mean of the inter-channel correlations is computed along with the global theta-band power and shown as a scatterplot, with one point per time window.. The distributions yield positive correlation.

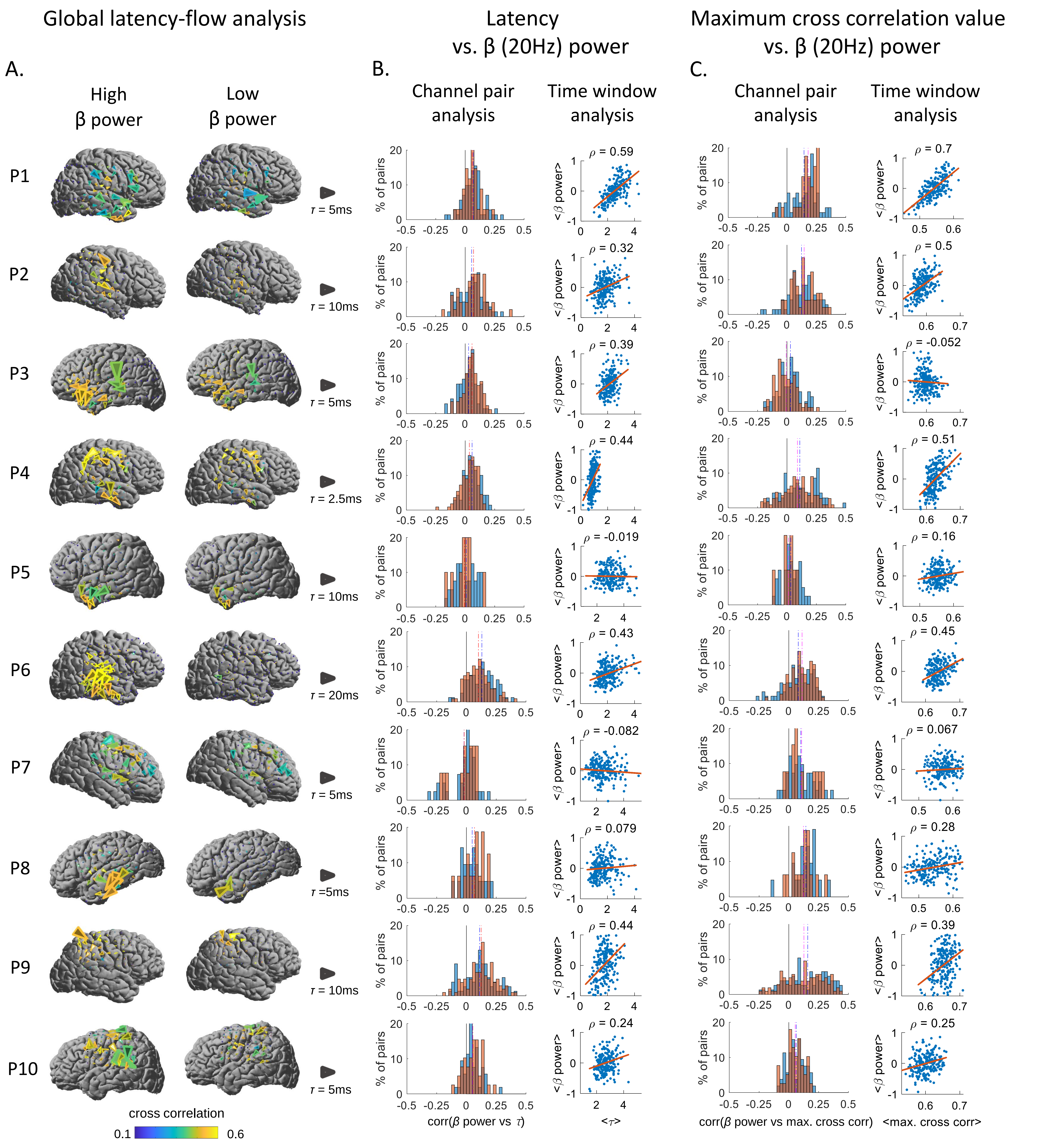


**Fig. S3. Latencies and coupling strengths increase with beta band power. (*A*) *Global latency-flow analysis*.** Brain surfaces illustrate latency flow maps averaged across time windows with high values (top 10% ) and low values (bottom 10%) of global beta-band power. Top 10% of the windows yielding highest beta power and bottom 10% of the windows yielding lowest beta power are chosen, and latency flows are computed across the chosen windows respectively. Yellower arrows denotes higher cross correlation. Larger size of arrows denotes longer latencies. The size of arrows on each flow map is scaled for readability: the grey arrow next to each participant's brain map indicates scale of latency flow arrows for that participant. (***B***) ***Time delay vs. beta power***. *Channel pair analysis*: For each pair of channels, the inter-channel latency and global beta power are correlated across time windows and the distribution of correlation values is shown in histograms. Red histograms represent nearest neighbor channel pairs and blue histograms represent next-nearest neighbor pairs. The distribution is positively skewed. *Time window analysis*: The mean latencies across pairs and mean global beta power for each time window are computed and shown as a scatterplot, with one point per time window. (***C***) ***Maximum cross correlation vs. beta power.*** *Channel pair analysis*: For each pair of channels, the inter-channel peak cross-correlation value is correlated across time windows with global beta power. The resulting correlation values are shown in the histogram. *Time window analysis*: For each time window, the spatial mean of the inter-channel correlations is computed along with the global beta-band power and shown as a scatterplot, with one point per time window.. The distributions yield positive correlation.

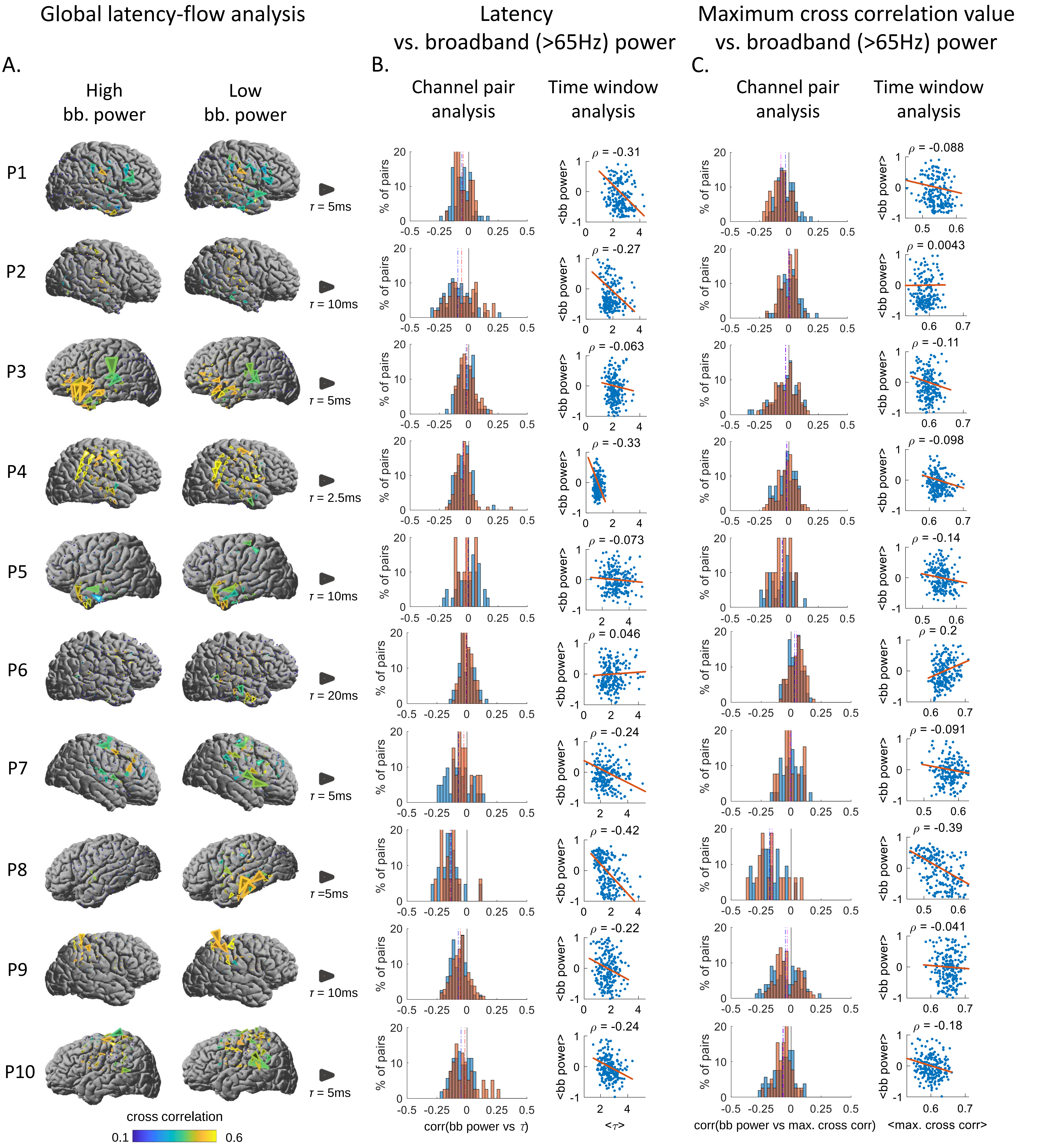


**Fig. S4. Latencies and coupling strengths decrease with broadbamd band power. (*A*) *Global latency-flow analysis*.** Brain surfaces illustrate latency flow maps averaged across time windows with high values (top 10% ) and low values (bottom 10%) of global broadbamd-band power. Top 10% of the windows yielding highest broadbamd power and bottom 10% of the windows yielding lowest broadbamd power are chosen, and latency flows are computed across the chosen windows respectively. Yellower arrows denotes higher cross correlation. Larger size of arrows denotes longer latencies. The size of arrows on each flow map is scaled for readability: the grey arrow next to each participant's brain map indicates scale of latency flow arrows for that participant. (***B***) ***Time delay vs. broadband power***. *Channel pair analysis*: For each pair of channels, the inter-channel latency and global broadbamd power are correlated across time windows and the distribution of correlation values is shown in histograms. Red histograms represent nearest neighbor channel pairs and blue histograms represent next-nearest neighbor pairs. The distribution is positively skewed. *Time window analysis*: The mean latencies across pairs and mean global broadbamd power for each time window are computed and shown as a scatterplot, with one point per time window. (***C***) ***Maximum cross correlation vs. broadband power.*** *Channel pair analysis*: For each pair of channels, the inter-channel peak cross-correlation value is correlated across time windows with  global broadbamd power. The resulting correlation values are shown in the histogram. *Time window analysis*: For each time window, the spatial mean of the inter-channel correlations is computed along with the global broadbamd-band power and shown as a scatterplot, with one point per time window.. The distributions yield positive correlation.

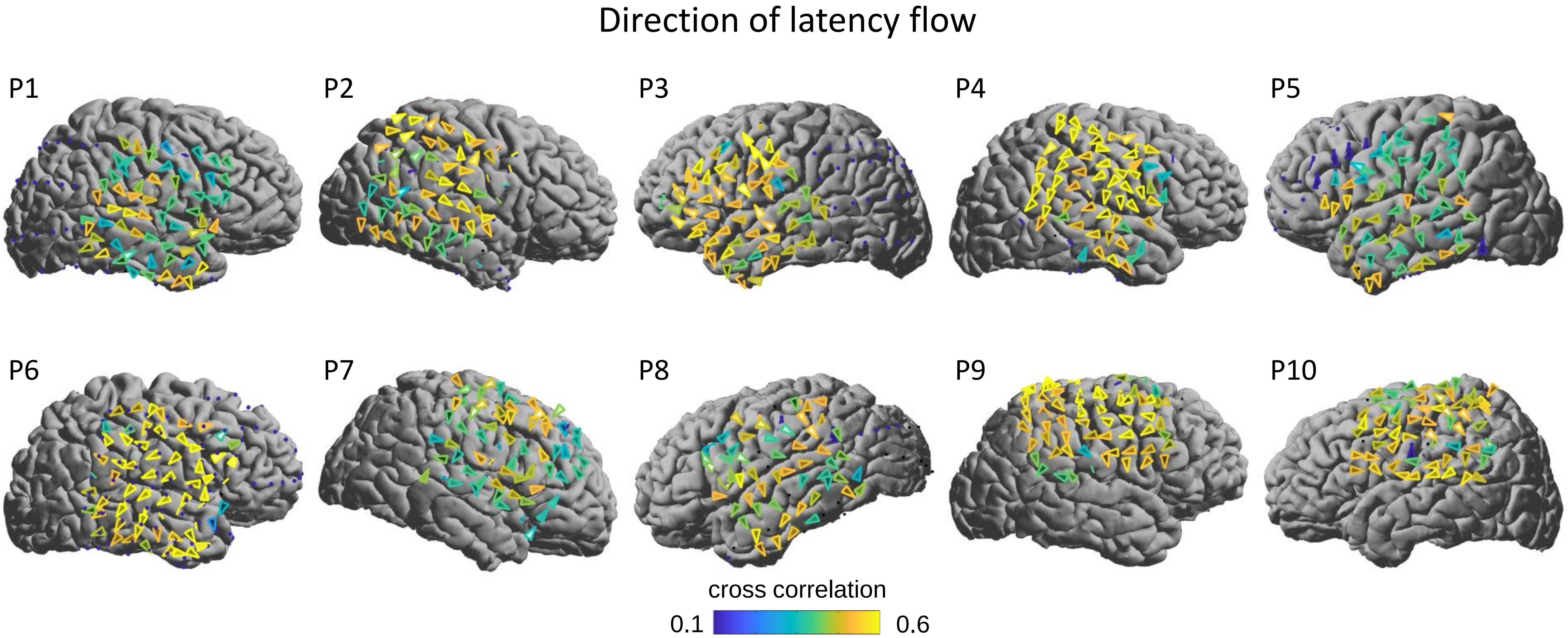


**Fig. S5**. **Latency flow.** Each figure shows the latency flow patterns averaged across time windows of elevated alpha band (10 Hz) global power. Time windows were sorted according to global alpha-band power, and latency flow was averaged across the top 10% of time windows (i.e. top 20 windows). Yellower color of arrows denotes higher cross correlation. In order to emphasize the direction of flow, all arrows are shown at the same size, and they are not weighted by latency.

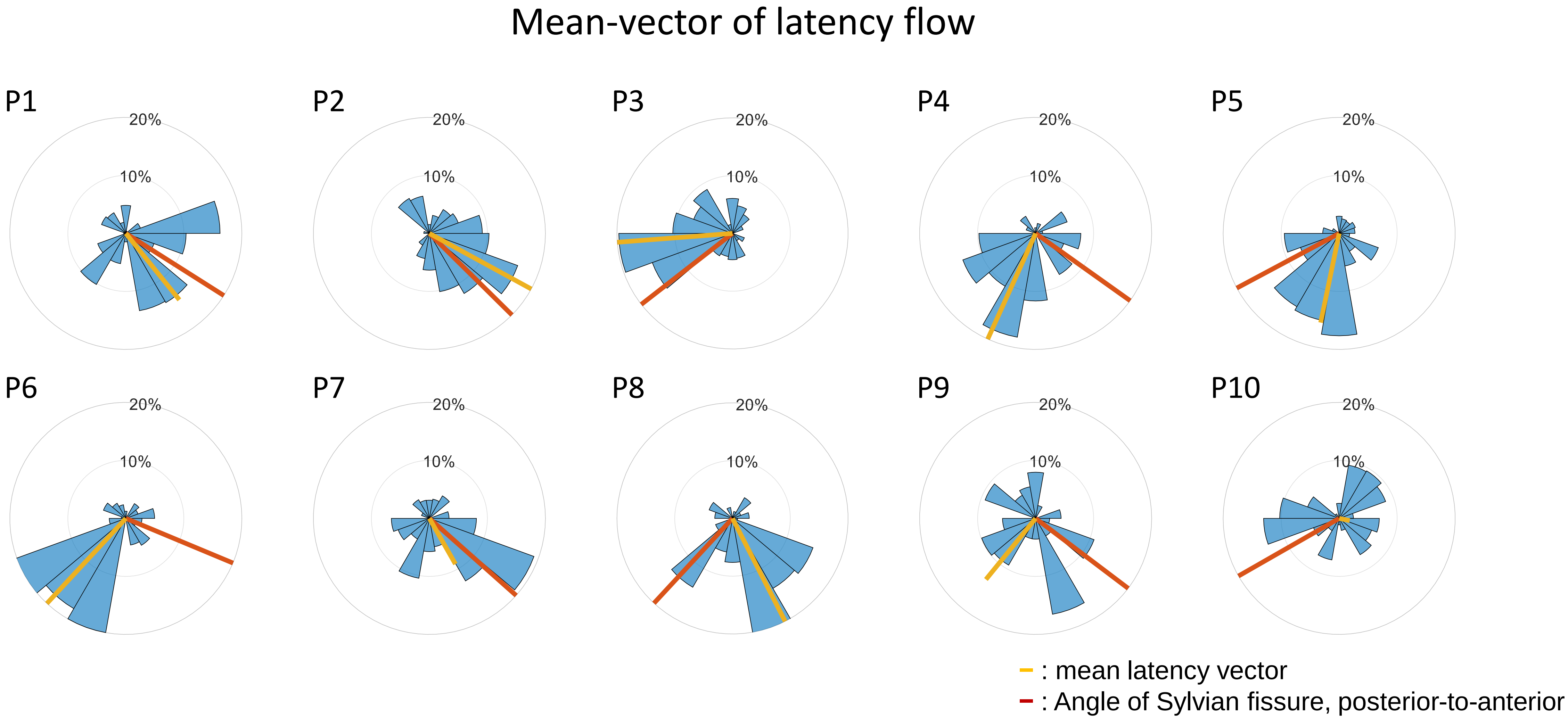


**Fig. S6. Mean latency flow vector relative to the Sylvian fissure.** Each figure shows the angle distribution of the latency flow vectors, and the mean-vector of latency flow (–) and the Sylvian Fissure orientation (–) for each participant. For 7 out of 10 participants (P1, P2, P3, P4, P5, P7, P8), the angle between the mean-vector and the Sylvian Fissure line is less than $0.5\pi$.

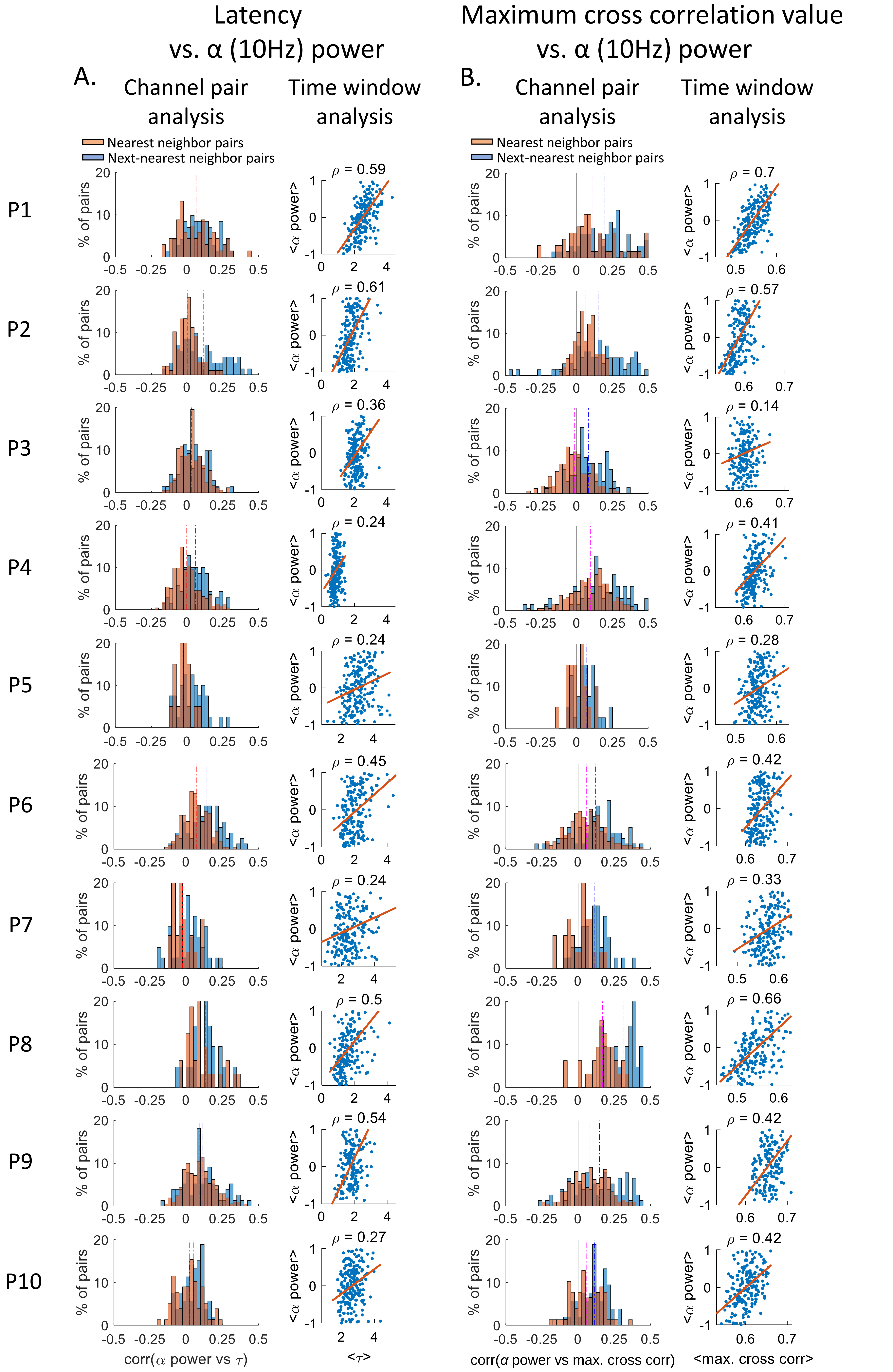


**Fig. S7. Latencies and coupling strengths increase with local alpha-band power.** (***A***) ***Time delay vs. alpha power.*** *Channel pair analysis*: For each pair of channels, the inter-channel latency and local alpha power are correlated across time windows and the distribution of correlation values is shown in histograms. Red histograms represent nearest neighbor channel pairs and blue histograms represent next-nearest neighbor pairs. The distribution is positively skewed. *Time window analysis*: The mean latencies across pairs and mean local alpha power for each time window are computed and shown as a scatterplot, with one point per time window. (***B***) ***Maximum cross correlation vs. alpha power.*** *Channel pair analysis*: For each pair of channels, the inter-channel peak cross-correlation value is correlated across time windows with local alpha power. The resulting correlation values are shown in the histogram. *Time window analysis*: For each time window, the spatial mean of the inter-channel correlations is computed along with the local alpha-band power and shown as a scatterplot, with one point per time window.. The distributions yield positive correlation.

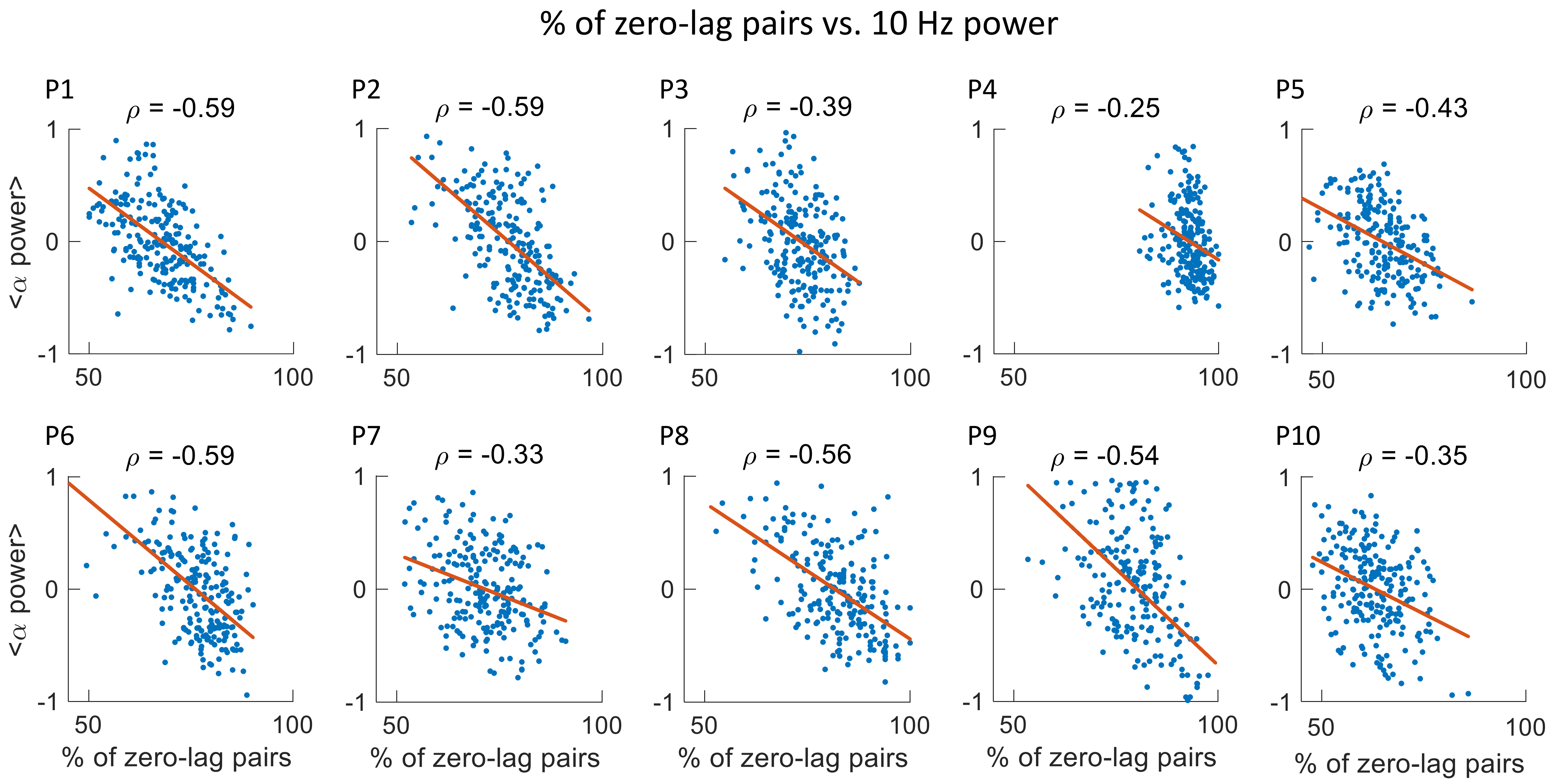


**Fig. S8. % of zero-lag pairs vs. *α* power.** Each figure shows % of zero-lag pairs (defined as latency in range [-2, 2] msec) vs. global alpha-band power, per each time window.

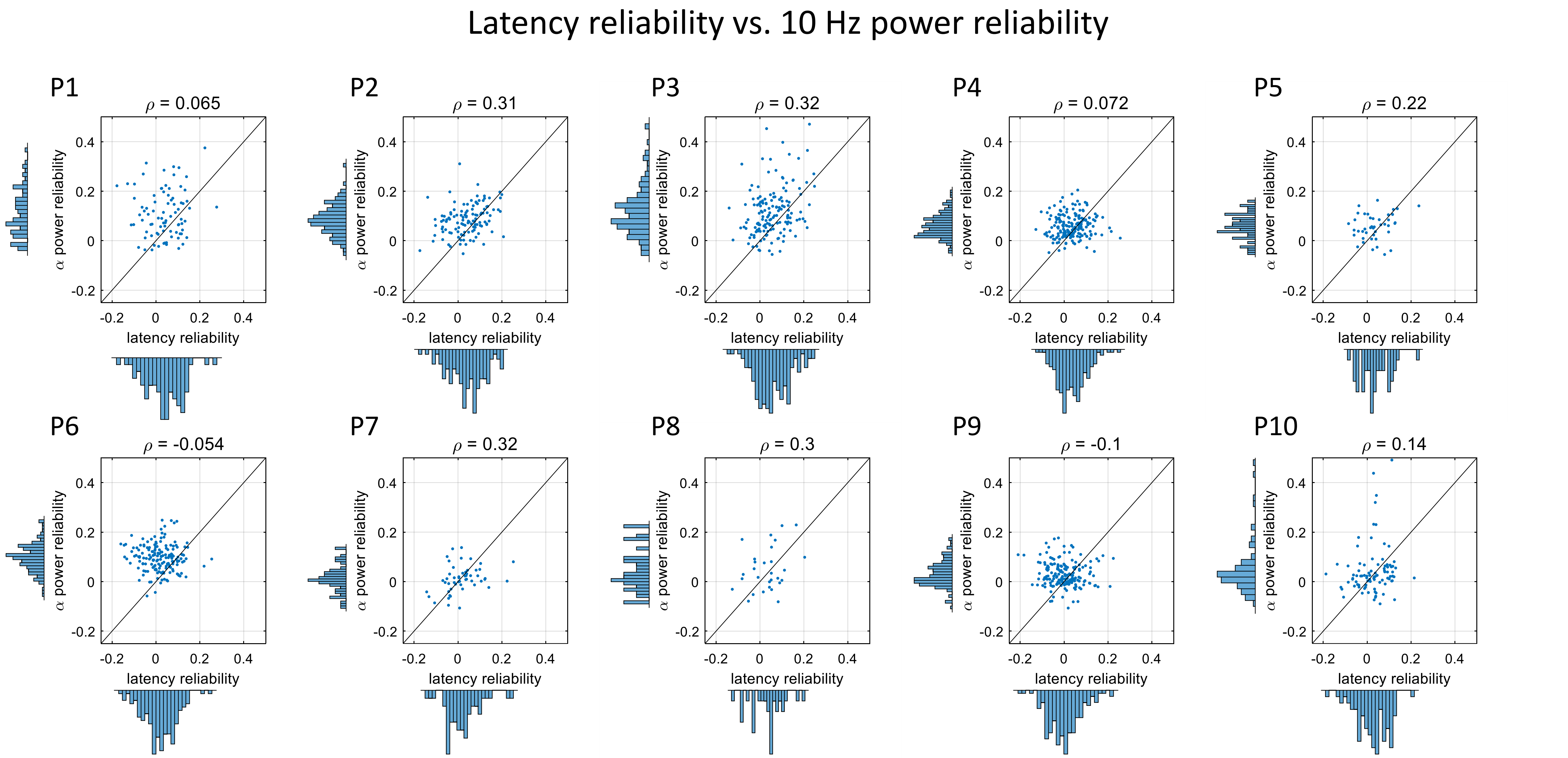


**Fig. S9. Latency reliability vs. *α* power reliability.** For each participant, latency reliability vs. alpha power reliability for each channel pair is shown.

### Latency reliability vs. 65+ Hz power reliability

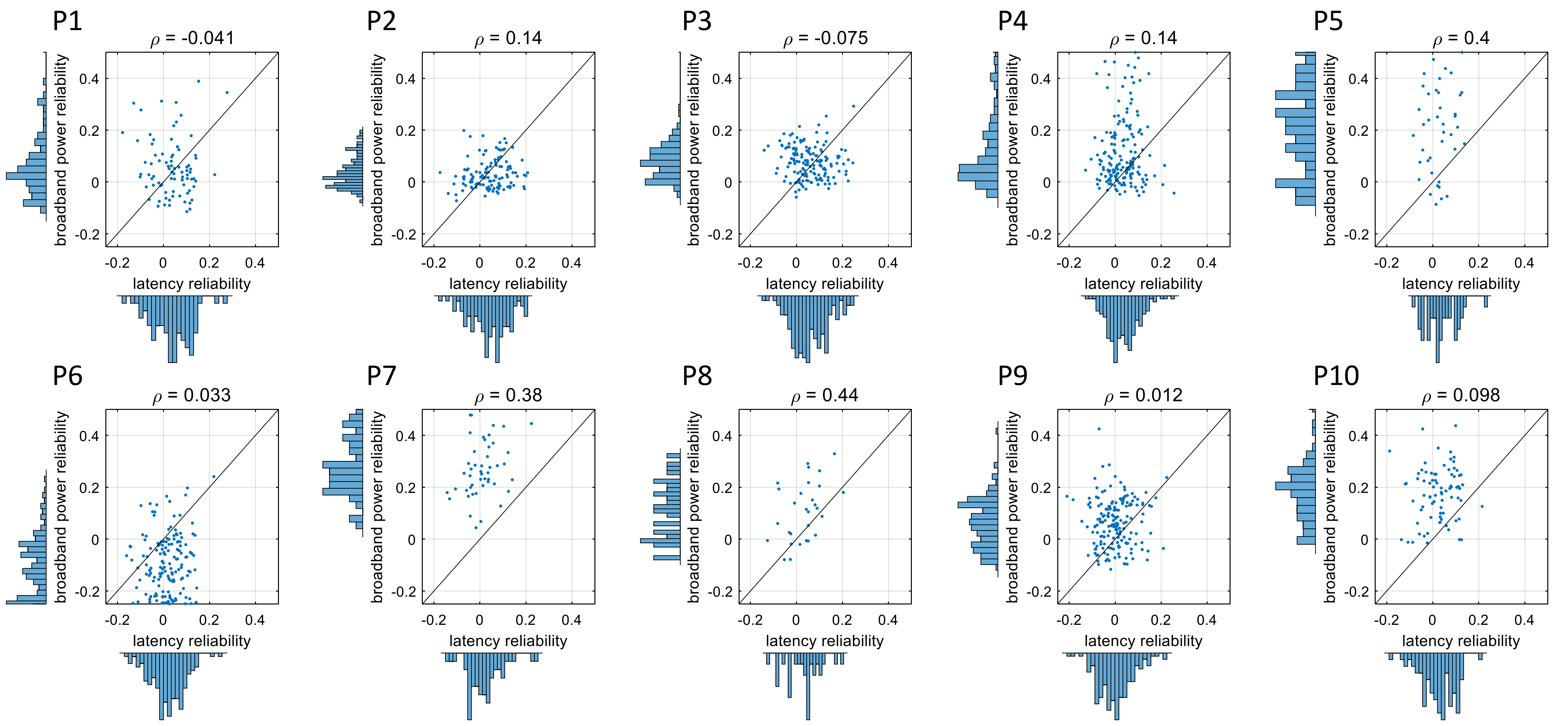


**Fig. S10. Latency reliability vs. broadband power reliability.** For each participant, latency reliability vs. broadband power reliability for each channel pair is shown.

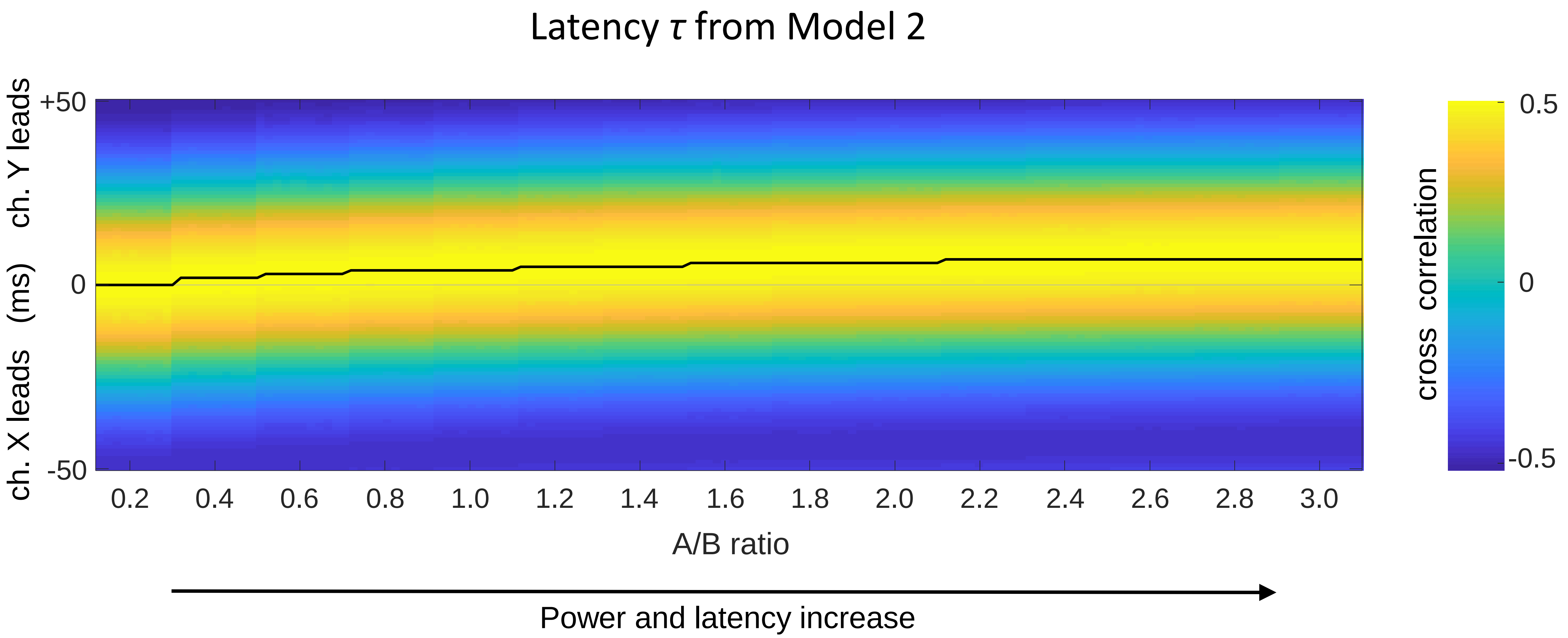


**Fig. S11. Measured latency *τ* from Model 2.** Cross correlation between two signals X and Y from Model 2 in the Supporting Text is shown. Each column of the colormap presents the cross-correlation from a model simulation using a different A/B ratio. Black line denotes the inter-electrode latency *τ*. As the ratio A/B increases, both the power of the signals and the latency *τ* between channel X and Y increases.
